# Extracellular Matrix Stiffness Alters TRPV4 Regulation in Chondrocytes

**DOI:** 10.1101/2021.09.14.460172

**Authors:** Nicholas Trompeter, Cindy J. Farino, Mallory Griffin, Ryan Skinner, Omar A. Banda, Jason P. Gleghorn, John. H Slater, Randall L. Duncan

**Affiliations:** Department of Biomedical Engineering, University of Delaware, Newark, DE; Department of Biology, University of Michigan-Flint, Flint, MI

**Keywords:** ECM Stiffness, TRPV4, Apoptosis, OA, Calcium

## Abstract

During the progression of osteoarthritis (OA), degradation of the extracellular matrix alters the biomechanical properties of cartilage, especially the compressive modulus. The mechanosensitive ion channel transient receptor potential vanilloid 4 (TRPV4) is required for chondrocyte mechanotransduction However, how OA-mediated cartilage degradation influences TRPV4 signaling remains unknown. To determine if ATDC5 cells alter TRPV4-mediated calcium signaling and cell phenotype in response to softer substrates, we created PEGDA-RGDS hydrogels with Young’s moduli that simulated healthy (~350 kPa), OA (~175 kPa) and severe OA (~35 kPa) tissue. We found that softer substrates reduced the influx of calcium through TRPV4 when challenging chondrocytes with hypotonic swelling (HTS). Chondrocyte apoptosis also increased on the OA and severe OA gels due to elevated basal [Ca^2+^]_i_, which is attenuated with pharmacological agonism of TRPV4. Pharmacological agonism of TRPV4 rescued the expression of aggrecan and TRPV4 in chondrocytes cultured on OA gels and enhanced the type II collagen (col2) expression in cells on the normal and OA gels. These data suggest that the biomechanical properties of degraded cartilage alter TRPV4-mediated mechanotransduction in chondrocytes. Given that TRPV4 reduced apoptosis and improved the chondrogenic capacity of cells, TRPV4 stimulation could provide a potential therapeutic target in patients with early to moderate OA.

## Introduction

As osteoarthritis (OA) progresses, altered biomechanical loading and extracellular matrix remodeling result in an overall decrease in the Young’s modulus of articular cartilage. Healthy knee cartilage has Young’s moduli between 0.4 – 2 MPa (Setton et al., 1999) and in severe OA the Young’s modulus can decrease up to an order of magnitude (Graham et al., 2018). Concomitantly, the tissue loses proteoglycans and becomes more permeable, potentiating the hydrodynamic and osmotic forces chondrocytes experience during OA (Hashimoto et al., 1998). Many studies have investigated how substrate elasticity and biomechanical loading influence proteoglycan and collagen production in chondrocytes embedded within hydrogels (Nicodemus et al., 2011; Nicodemus and Bryant, 2008; Nims et al., 2021; O’Conor et al., 2014; Schuh et al., 2012). However, few studies have investigated the role of specific mechanotransduction pathways, specifically mechanosensitive ion channels, during the degradation of articular cartilage.

A key mechanosensor within chondrocytes is the non-selective cation channel Transient Receptor Potential Vanilloid 4 (TRPV4). TRPV4 mediates the response of chondrocytes to dynamic mechanical load and osmotic tension (O’Conor et al., 2014) that is critical for chondrocyte homeostasis (Urban et al., 1993). Additionally, stimulation of TRPV4 reduces inflammation by inhibiting interleukin-1β mediated cartilage degradation (Fu et al., 2021). Pharmacological agonism of the TRPV4 channel has also shown therapeutic promise in promoting chondrogenic differentiation and inhibiting cartilage degradation during post-traumatic OA (Atobe et al., 2019). These studies underscore TRPV4 as a regulator of chondrocyte mechanotransduction and as a viable therapeutic target for treating OA. However, how changes in the extracellular environment during OA alter TRPV4 mediated mechanotransduction and its influence on the phenotype of chondrocytes is unknown.

Polyethylene glycol (PEG) hydrogels have been utilized for cartilage engineering due to 1) their biocompatibility, 2) the ability to tether biomimetic peptides/proteins and drugs to the hydrogels, and 3) their tunable mechanical properties (Bryant et al., 2005; Elisseeff et al., 1999). Unfortunately, chondrocytes embedded in PEG hydrogels with compressive moduli similar to healthy articular cartilage have worse biomechanical properties and a stunted chondrogenic phenotype when compared to hydrogels with compressive moduli similar to OA cartilage (Bryant et al., 2005, 2004; Nicodemus and Bryant, 2008). PEG hydrogel models have great potential as engineered cartilage constructs because chondrocytes within these hydrogels deposit collagen and proteoglycan, improving their biomechanical properties over a matter of days/weeks (Galarraga et al., 2019). However, their capacity to bestow a chondrogenic phenotypic to chondrocytes and mesenchymal stem cells limits their utility to investigate how ECM remodeling during OA influences TRPV4-mediated mechanotransduction.

By seeding chondrocytes on the surface of PEG hydrogels, we sought to determine if alterations to substrate elasticity influences TRPV4 signaling. Furthermore, we postulated that pharmacological stimulation of TRPV4 may enhance the anabolic phenotype of cells grown on hydrogels that recapitulate the Young’s moduli of OA and severe OA tissues.

## Methods

### Preparation of tunable PEGDA-RGDS hydrogels

(**Fig. 1**) PEG-Diacrylate (PEGDA) was synthesized as previously reported (Banda et al., 2019). Briefly, poly(ethylene-glycol) (3.4kDa, Sigma-Aldrich) was reacted with acryloyl chloride (AC, Sigma-Aldrich) and triethylamine (Sigma-Aldrich) in dichloromethane for 24 hrs. PEGDA was then purified by precipitation in 0°C diethyl ether. Monoacrylate PEG-succinimidyl ester (3400 Da SVA, Laysan Bio) was mixed with the integrin-ligating peptide RGDS (Bachem) in DMSO with N,N-diisopropylethylamine for 24 h. A 5.2%, 9.8%, and 14.1% (w/v) of PEGDA was added to 10 mM, 4.37 mM, and 3.97mM PEG-RGDS, respectively, in Hank’s Balanced Salt Solution (HBSS, Corning) and mixed with 3 mg/ml of lithium phenyl-2,4,6-trimethylbenzoylphosphinate. The solutions were pipetted on individual perfluoroalkoxy coated glass slides (McMaster) between 150 μm polydimethylsiloxane (PDMS) spacers before polymerizing the gels with UV light (Blak-Ray flood UV lamp, intensity of 10 mW/cm^2^ at 365 nm) for 60 s. This created flat 150 μm thick PEGDA-RGDS gels that were ~0.325 cm in radius.

**Figure 1.**
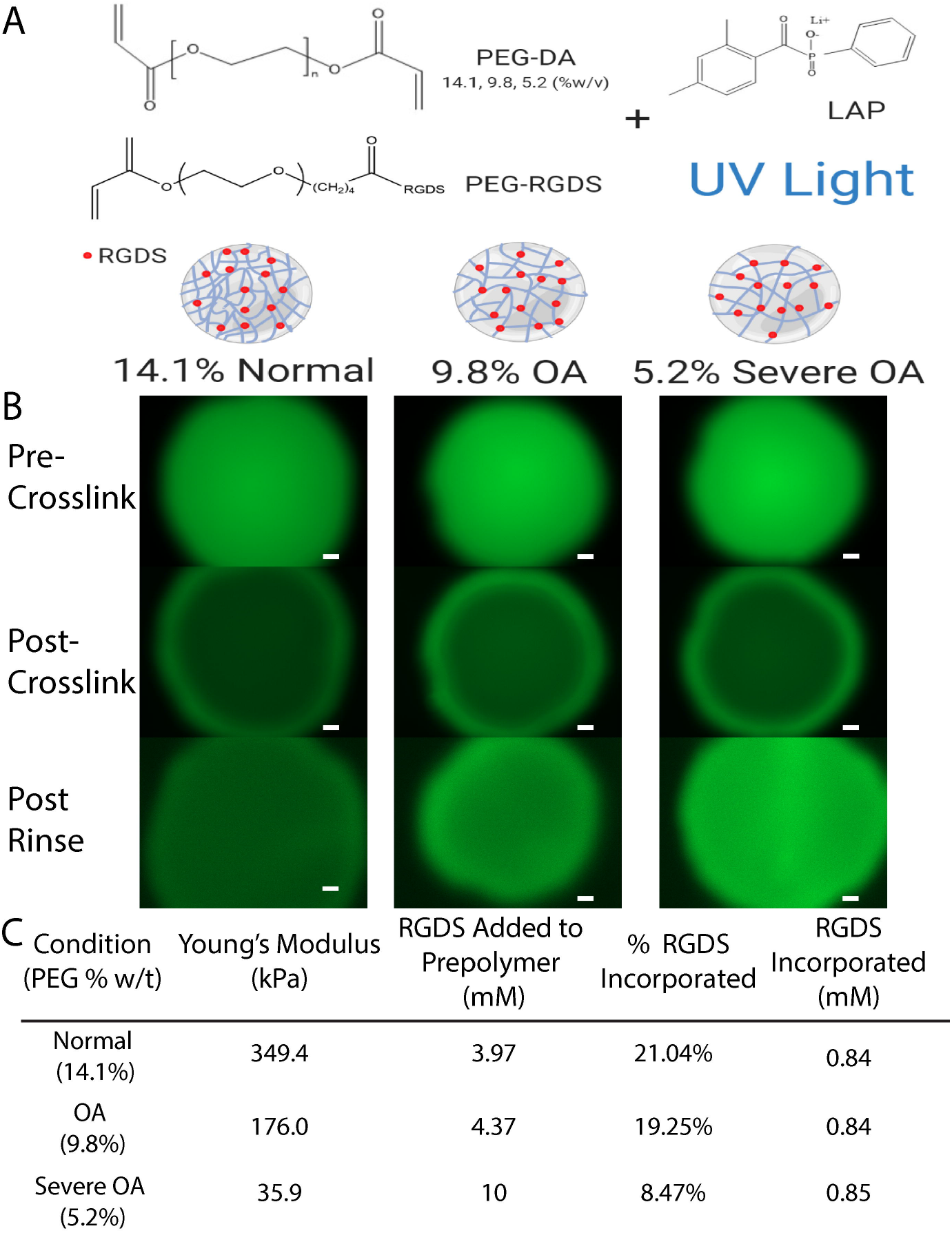
A) Illustration of the strategy used to generate tunable PEG-RGDS hydrogels of normal (14% PEG) OA (9.8% PEG) and severe OA (5.2% PEG) with equal incorporation of RGDS across the platform. C) The fluorescent intensity of PEG-RGDS-488 in the normal, OA, and severe OA gels before crosslinking, post-crosslinking with UV light, and after washing the gels for 24 hrs. Scale bars = 200 μm. C) Properties of the PEG-RGDS hydrogels that were utilized to recreate normal, OA, and severe OA cartilage. Made in BioRender Software

To normalize adhesion ligand density of the RGDS, PEG-RGDS incorporation efficiency was quantified for gels composed of 5.2%, 9.8%, and 14.1% PEGDA. Fluorescence analysis was implemented using a fluorophore analog of RGDS to visualize peptide uniformity as previously reported (Pradhan et al., 2019). Briefly, a PEG-RGDS-Alexa Fluor 488 (PEG-RGDS-488) was synthesized in a similar manner as PEG-RGDS, with Alexa Fluor 488 succinimidyl ester (Thermo Fisher) dissolved in the reaction mixture of PEG-SVA and RGDS. Gels made up of 5.2%, 9.8%, and 14.1% PEGDA, with 0.5 mM PEG-RGDS-488, and 9.5 mM PEG-RGDS were prepared. Pre-polymer solutions were imaged before photopolymerization to account for bleaching, immediately after polymerization, and after overnight rinsing in HBSS. Intensity measurements before and after rinsing were used to determine conjugation efficiency as follows:

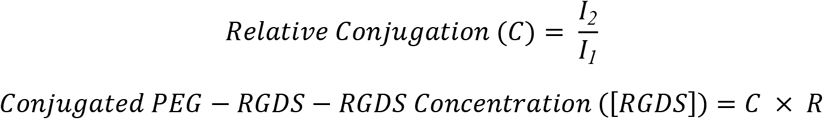

Where I_1_ is the fluorescence post-crosslinking, I_2_ is the fluorescence after washing, and R is the initial prepolymer concentration of PEG-RGDS in solution. The incorporation of RGDS into the 5.2% PEGDA hydrogels was used to equalize the concentration of RGDS into the 9.8% and 14.1% PEGDA hydrogels.

### Mechanical testing

To formulate PEG hydrogels with Young’s moduli of ~35, ~175, and ~350 kPa we first determined the Young’s moduli of hydrogels of 6.2%, 13.5%, and 20.1% (w/v) PEGDA. Hydrogels 8 mm in diameter and 1.3 mm thick had constant strain applied under unconfined compression (0.18 mm/min, Instron 5934, 100N). The strain at 20% was utilized to determine the Young’s moduli of these gels. This data fit to a second order polynomial to determine the percentages of PEG that theoretically recapitulate hydrogels of the desired Young’s moduli. Hydrogels of 5.2%, 9.8%, and 14.1 % (w/v) PEGDA were tested as above to confirm their Young’s moduli (**Table 1**).

**Table 1.** Ca^2−^ kinetics in response to HTS and GSK219 on PEG-RGDS hydrogels.

|  | Normal Control | Normal GSK219 | OA Control | OA GSK219 | Severe OA Control | Severe OA GSK219 |
| --- | --- | --- | --- | --- | --- | --- |
| Baseline Prior to HTS | 24.88 ± 0.89<sup>A,B</sup> | 23.51 ± 0.88<sup>A</sup> | 32.05 ± 1.47<sup>C</sup> | 32.24 ± 1.96<sup>B,C</sup> | 56.92 ± 2.42<sup>D</sup> | 55.22 ± 2.45<sup>D</sup> |
| Cells Responding (%) | 82.10 ± 8.42<sup>A</sup> | 67.46 ± 8.35<sup>A,B</sup> | 54.94 ± 9.42<sup>A,C</sup> | 22.22 ± 15.32<sup>C</sup> | 30.86 ± 5.25<sup>C</sup> | 31.25 ± 7.42<sup>B,C</sup> |
| Time to Peak after HTS (s) | 19.09 ± 1.16<sup>A</sup> | 25.23 ± 1.40<sup>B</sup> | 19.86 ± 1.34<sup>A,B</sup> | 18.51 ± 2.94<sup>A,B</sup> | 19.61 ± 1.79<sup>A,B</sup> | 20.94 ± 1.87<sup>A,B</sup> |

Mean values ± SEM. Conditions with the same superscript are not significantly different from one another. p-value < 0.05.

### Calcium imaging

Calcium imaging experiments were conducted as described previously, (Hurd et al., 2015; Trompeter et al., 2021) with slight modifications to the protocols. Briefly, the chondrogenic cell line ATDC5 (passages 13 – 23), were seeded on the surface of normal (~350 kPa), OA (~175 kPa), and severely OA (~35 kPa) hydrogels at 21,000 cells/cm^2^ for hypotonic swelling (HTS) experiments, and 15,000 cells/cm^2^ for calcium oscillation studies. Cells attached for 1 h at 37°C before adding DMEM/Ham’s F12 media (Corning) supplemented with 10% FBS (Atlas Biologicals), and 1% penicillin/streptomycin (HyClone). Cells were cultured for 48 h prior to serum starvation in 0.2% FBS for 16 hr. For calcium oscillation experiments, cells were treated with 30 nM of TRPV4 agonist GSK1016790A (GSK101, MilliporeSigma) for 28 hrs before serum starvation. For HTS experiments, cells were washed twice in HBSS for 10 minutes and loaded with Fluo-4 AM (Invitrogen) for 15 minutes. Cells were then washed in HBSS, and recovered in HBSS or 100 nM GSK2193874 (GSK219, MilliporeSigma), a TRPV4 specific antagonist, at RT for 15 minutes. Cells were imaged on a 5 Live DUO Highspeed Confocal Microscope (Zeiss). A 1-minute baseline of fluorescence was taken before challenging the cells with 50% HTS. For calcium oscillation experiments, cells were loaded with 1 μM of Fluo-8 AM (MilliporeSigma) in 0.2% serum media for 30 minutes at 37°C, washed with 37°C HBSS, and recovered in 37°C HBSS. Calcium imaging was conducted at 37°C using a Zeiss Axio Observer Z1 inverted microscope. The peak, baseline, percent responding, percent stable, and the number of peaks per cell were analyzed using a custom MATLAB code. A cell was responding if a peak was 2.5% over baseline fluorescence. This threshold was determined by analyzing calcium traces from one set of normal gels. A cell was considered unstable if the baseline [Ca^2+^]_i_ decreased by more than 40% or increased by more than 20% (**Supplemental Fig 3).** Thresholds were chosen based upon photobleaching of the sample and the fluorescence of cells on normal gels, where less than 1% of cells were unstable.

### Apoptosis assays

Cells were seeded as above and were cultured for 48 hr. A subset of gels received 30 nM of GSK101 in growth media for 28 hr. After serum starvation, cells were incubated at 37°C for 30 minutes in 5 μM propidium iodide (Alfa Aesar) and 1:1500 Hoechst 33342 (Molecular Probes) in serum reduced media. Dead cells were counted using ImageJ. Apoptosis was confirmed using the CellEvent™ Caspase-3/7 Green Detection Reagent following the manufacturer’s recommendations. A 5 μM CellEvent™ Caspase-3/7 Green Detection Reagent and Hoechst solution were added to cells and incubated at 37°C for 1 hr. Cells were imaged using a Zeiss AxioObserver Z1 microscope. The percentage of Caspase 3/7 positive cells was quantified using a custom MATLAB code generously provided by Dr. Michael David.

### Immunofluorescence

Cells were seeded at 15,000 cells/cm^2^. A subset of cells received 30 nM GSK101 for 28 h. Cells were fixed in 4% paraformaldehyde, blocked for 1 hr in PBS containing 1% BSA/0.2% Fish Gelatin (Fisher Scientific), and probed for membrane TRPV4 using 1:200 rabbit anti-TRPV4 (Alomone Labs, ACC-124) overnight at 4°C. Phalloidin-iFluor 647 Reagent (Abcam), Hoechst, and goat anti-rabbit Dylight 550 antibody (1:1000, Life Technologies) were applied for 1 hr at RT. Confocal microscopy was performed on a Zeiss LSM 800. TRPV4 fluorescence was analyzed using ImageJ.

### qPCR

RNA was extracted from cells seeded on hydrogels using the ISOLATE II RNA Mini Kit (Bioline) following the manufacturer’s instructions. Individual Samples were run in duplicate using Taqman qPCR probes (Thermo Fisher) and the SensiFAST™ SYBR® No-ROX Kit (Bioline), following the manufacturer’s instructions. Using a LightCycler 96 system (Roche), expression of *TRPV4* (Mm00499025)*, Sox-9* (Mm00448840)*, Aggrecan* (Mm00545794), *Col2a* (Mm01309565), *Col1a1* (Mm00801666), and *MMP-13* (Mm00439491) were quantified using *Rplp0* (Mm00725448) as the internal control. The ΔΔCt values were normalized to the normal control for each gene and statistical analysis was completed using the normalized ΔΔCt.

### Western Blotting

To obtain sufficient protein from cells for western analysis, 40 μl hydrogels were used. ATDC5 cells were seeded on top of hydrogels at 7500 cells/cm^2^. Cells were lysed with RIPA Buffer supplemented with protease inhibitors and PMSF (MilliporeSigma). Protein concentrations were quantified using a Pierce BCA Kit (Thermo Fisher) before loading 6 μg of protein on Bolt Bis-Tris gels (Life Technologies), transferring to 0.2 mm nitrocellulose membranes (BioRad), and blocking overnight in 5% milk in tris-buffered saline with 0.1% Tween-20. TRPV4 and GAPDH were probed with rabbit anti-TRPV4 (1:500, Alomone Labs) and rabbit anti-GAPDH (1:5,000, Proteintech) overnight.

Membranes were then probed with goat anti-rabbit HRP (1:10,000, Licor) before being developed with SuperSignal West Femto (GE Life Sciences) and imaged on an iBright1500 (ThermoFisher).

### Statistical Analysis

All experiments were completed as both biological and technical triplicates. Significance between groups was determined using either one-way or two-way analysis of variance (ANOVA) with a Tukey post hoc test to determine a significance when multiple comparisons in the study were made with a p value <0.05 considered significant. All statistical analyses were completed using Origin software (Origin Lab).

## Results

PEGDA gels of 14.1%, 9.8%, and 5.2% (w/v) were used to capture the progression from healthy cartilage (~350 kPa) to severely OA cartilage (~35 kPa) **(Fig 1A.)**. To avoid confounding results based upon ligand density (Schuh et al., 2012; Villanueva et al., 2009), RGDS incorporation efficiency was analyzed and equalized across the three hydrogel formulations **(Fig 1B and C)**.

The intracellular calcium ([Ca^2+^]_i_) response of cells to HTS was analyzed across the three hydrogel stiffnesses; normal, OA, and severely OA. As gel stiffness decreased, the response to HTS also decreased **(Fig 2A.)**. Inhibition of TRPV4 with 100nM GSK219 attenuated the peak [Ca^2+^]_i_ and total Ca^2+^ influx in cells on the normal and OA gels, but not on the severe OA gels **(Fig 2B and C)**. When analyzing other parameters of Ca^2+^ kinetics, we observed that substrate elasticity influences basal calcium and the number of cells responding to HTS, but had no effect on the time to peak after HTS (Table 1). As stiffness of the gels decreases, the basal [Ca^2+^]_i_ increases; however, the number of cells responding concomitantly decreases **(Table 1)**.

**Figure 2.**
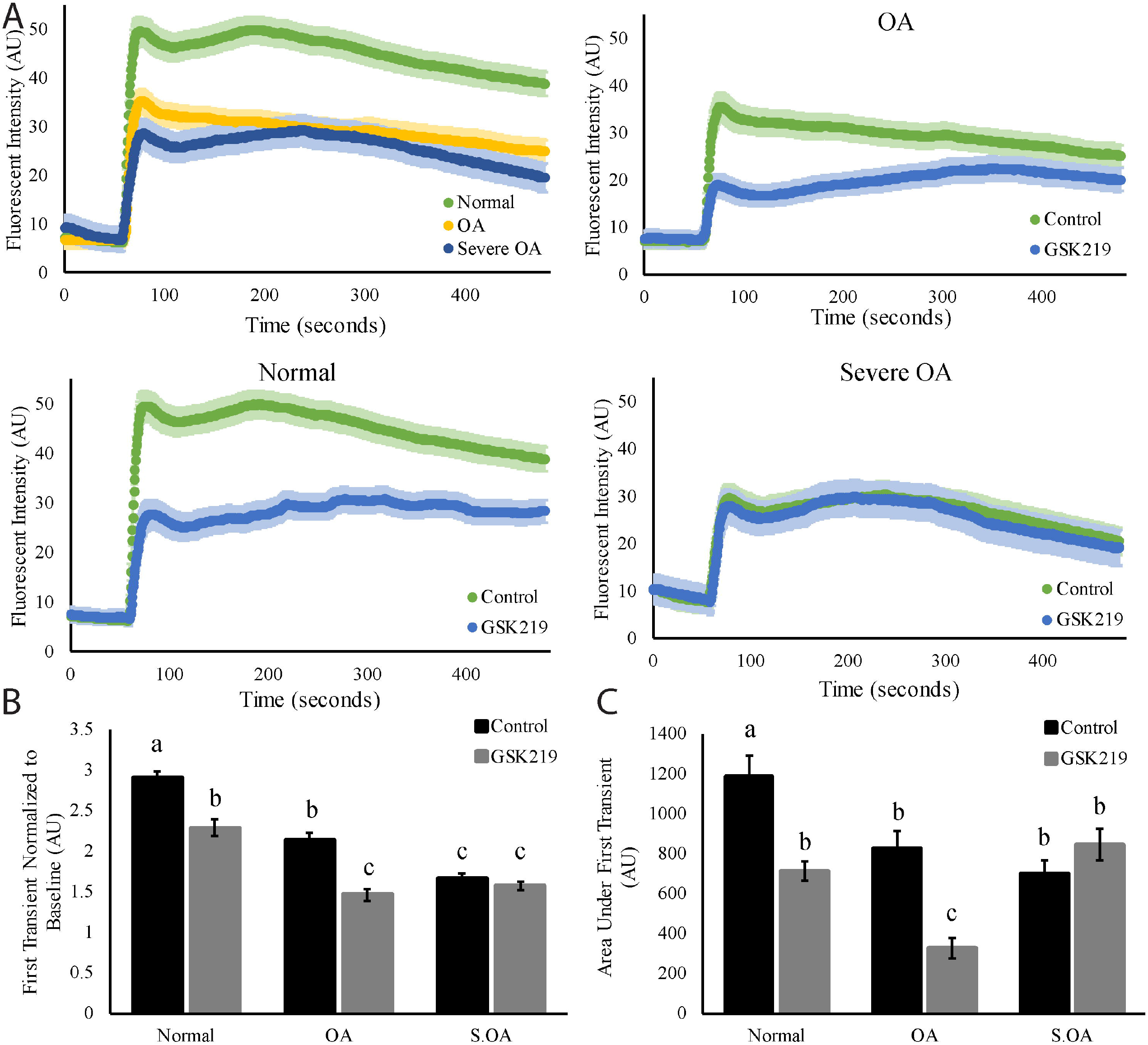
A) Average [Ca^2+^]_i_ traces ± SEM of ATDC5 cells on normal, OA, and Severe OA gels in response to 50% HTS. Upper left panel shows control conditions for normal (green), OA (blue), and Severe OA (yellow). All other panels compare control (green) and 100 nM GSK219(blue) treatments for each condition. All traces had normalized baselines to highlight the differences in the response to HTS. Actual baseline for each condition and treatment are found in Table 1. * indicates significant difference in first transient and area under the curve of the first transient (total [Ca^2+^]_i_) when compared to the control conditions. B) First transient (peak of [Ca^2+^]_i_ for normal, OA, and Severe OA conditions for control (black) and 100 nM GSK219 (gray) when normalized to the baseline during HTS. C) Area under first transient/peak is shown for control (black) and 100 nM GSK219 (gray) treatments to measure total [Ca^2+^]_i_ in response to HTS. For B) and C). Data with the same letter indicates that the conditions and/or treatments are not significantly different from one another and bars with different letters are significantly different from one another. Error bars are ± SEM, p-value <0.05.

When cells were challenged with HTS, there was a significant decrease in [Ca^2+^]_i_ influx in cells on the severely OA gels, which we attributed to a loss of cell membrane integrity due to rupture of a subset of the cell population **(Supplemental Movie 1)**. When evaluating cell viability, chondrocytes on the normal gels had significantly fewer apoptotic cells compared to chondrocytes on the OA and severe OA gels **(Fig 3A)**. Activating TRPV4 with 30 nM of GSK101 for 28 hr, significantly decreased the number of apoptotic cells when compared to the untreated severe OA group **(Fig 3A)**. Caspase 3/7 staining confirmed the increase in cell apoptosis on the OA and severe OA gels **(Fig 3B)** but TRPV4 stimulation failed to decrease the number of apoptotic cells on the severe OA gels.

**Figure 3.**
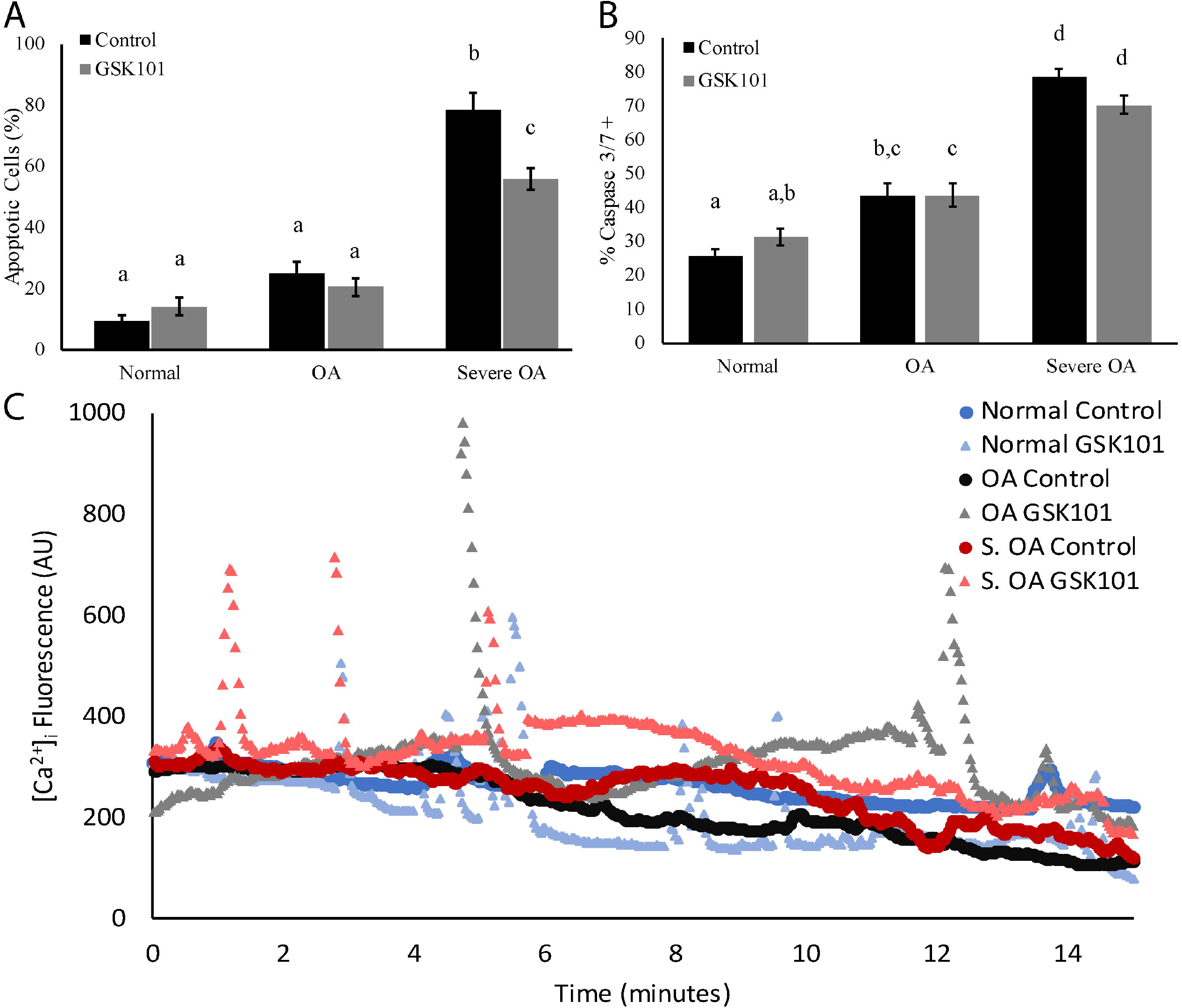
A) The percentage of ATDC5 cells positive for propidium iodide on normal, OA, and Severe OA gels during control (black) or 28 hr of 30 nM GSK101 (gray) treatments. Apoptosis/PI staining increases as the gels become softer. B) The percentage of ATDC5 cells positive for the apoptosis markers of Caspase 3/7 on normal, OA, and Severe OA gels during control (black) or 28 hr of 30 nM GSK101 (gray) treatments. A similar trend in Caspase 3/7 positive cells as seen in the PI assay. C) Representative traces of ATDC5 [Ca^2+^]_i_ for normal (dark blue, circles), OA (dark yellow, circles), severe OA (dark purple, circles) normal GSK101 (light blue, triangles), OA GSK101 (light yellow, triangles), and severe OA GSK101 (light purple, triangles) at 37°C. The baselines of all traces were decreased to better display the data. Data with the same letter indicates that the conditions and/or treatments are not significantly different from one another and bars with different letters are significantly different from one another. Error bars are ± SEM, p-value <0.05.

To determine why chondrocyte apoptosis increased on the OA and severe OA gels, we analyzed spontaneous calcium oscillations **(Fig. 3C)** that have been shown to promote the biomechanical properties of cartilage (Zhou et al., 2015). Stimulation of TRPV4 significantly increased the number of peaks per responding cell and the initial baseline [Ca^2+^]_i_ in the normal and OA conditions when compared to controls **(Table 2)**. Additionally, in control conditions, basal calcium was increased in the severe OA treatment group when compared to the normal and OA control groups (S.OA vs. normal and OA: 1484.20 ± 66.46 vs. 600.86 ± 13.81 and 679.73 ± 23.15) (Table 2). The magnitude of the peaks differed significantly between the severe OA control and the normal gels treated with GSK101 **(Table 2)**. However, chondrocytes on the severe OA gels had significantly more unstable cells compared to the other conditions as noted by drastic declines in [Ca^2+^]_i_ or non-uniform baseline [Ca^2+^]_i_.

**Table 2.**
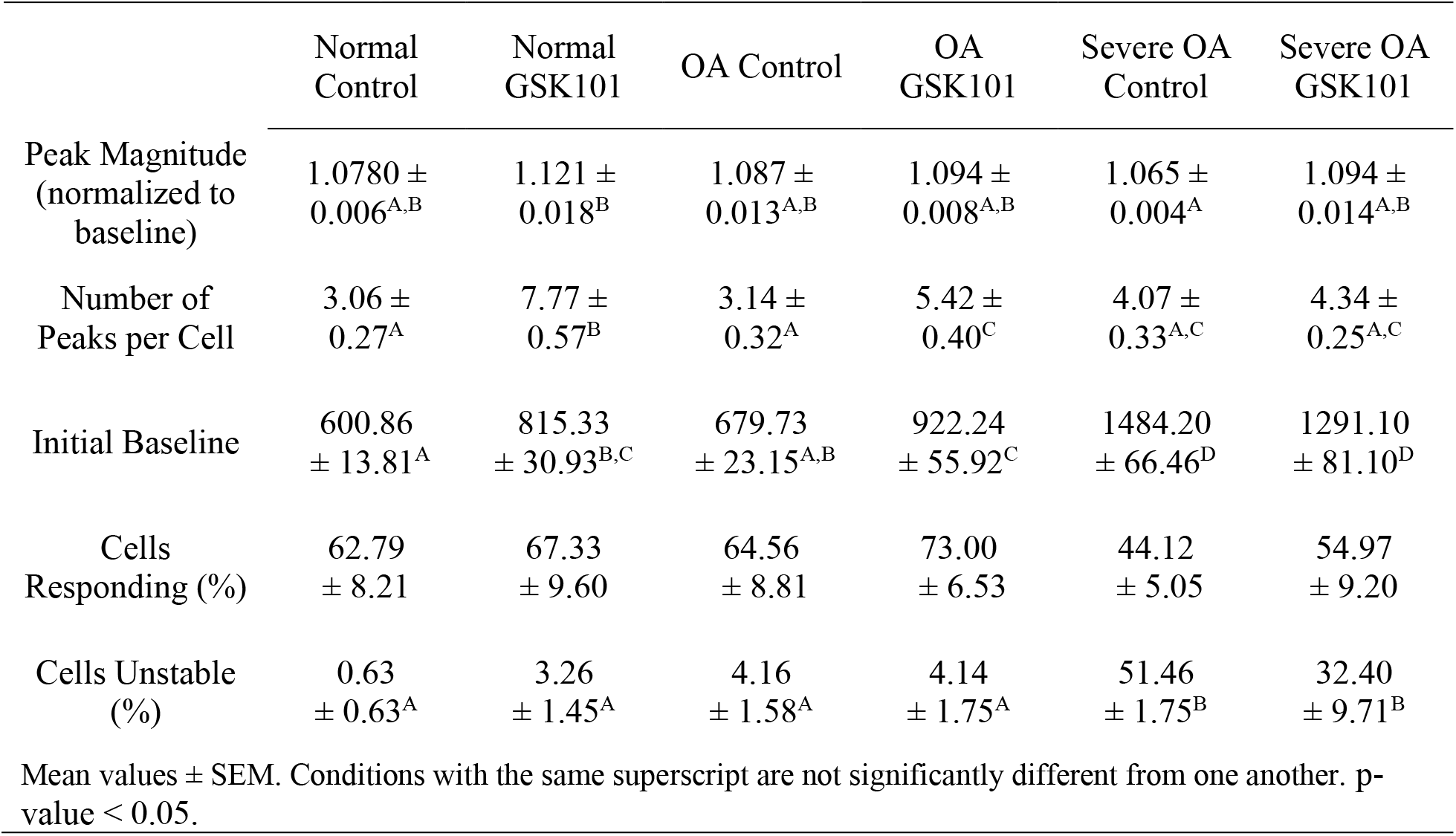
Ca^2+^ kinetics during calcium oscillations with GSK101 pretreatment on PEG-RGDS hydrogels.

We also sought to determine if changes in substrate elasticity altered TRPV4 expression. Cells on all three gel stiffnesses exhibited immunostaining of TRPV4 localized to the membrane **(Fig 4A)**. Expression of membrane bound TRPV4 was only significantly different between the normal gels treated with GSK101 and the severe OA controls **(Fig 4B and Supplemental Figure 4)**. Spatial distribution of the cells did not appear to differ between groups (data not shown).

**Figure 4.**
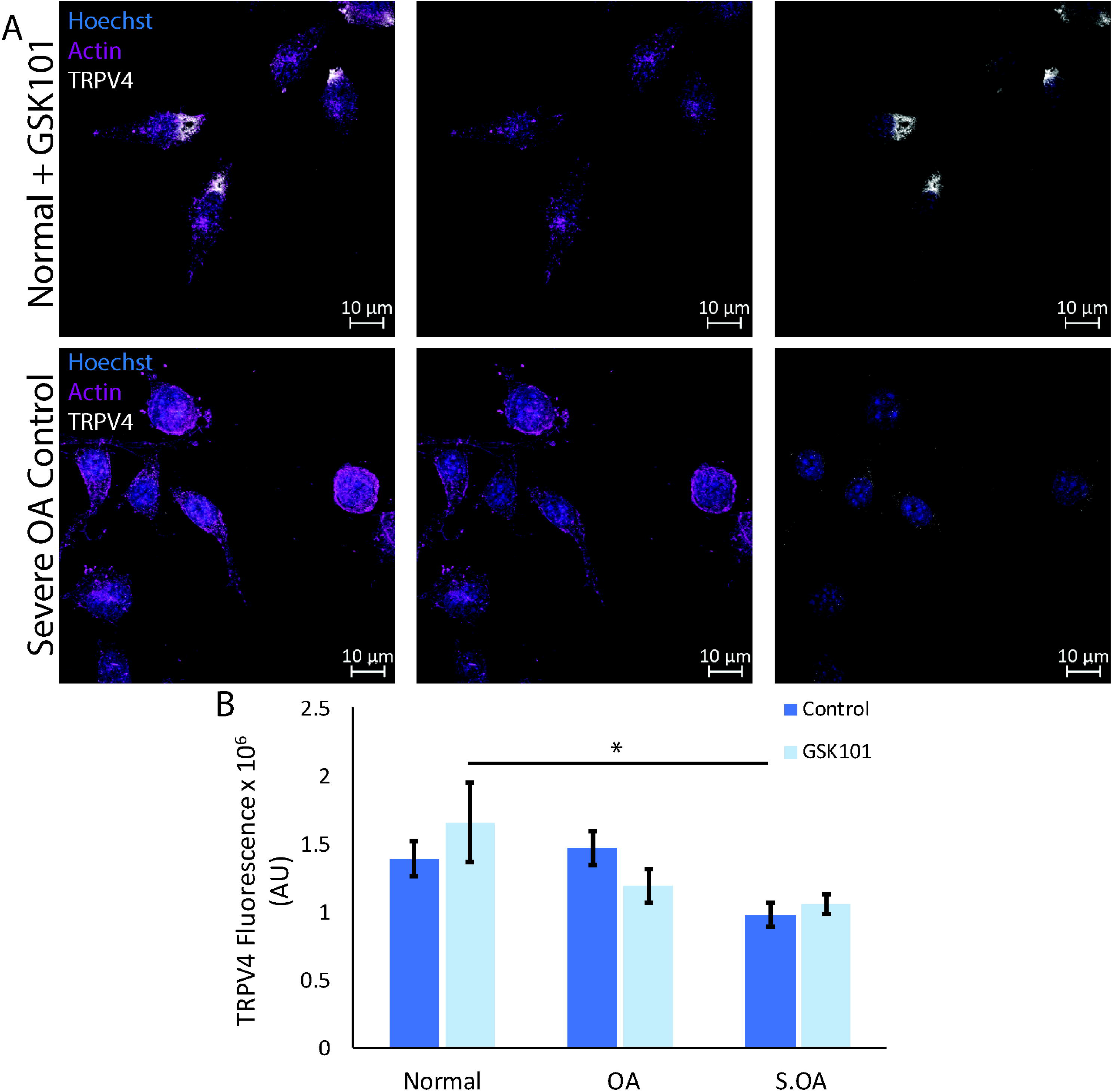
A) Immunofluorescent images of ATDC5 cells on normal gels (upper panel) treated with 30nM GSK101 for 28 hr and severe OA gels with control treatment (lower panel) stained for the nucleus (Hoechst, blue), f-actin (purple), and TRPV4 (white). The left panel shows the composite images of all stains (scale bar =10 μm). The middle panel shows nuclear and f-actin staining, and the right panel shows TRPV4 and nuclear staining. B) TRPV4 fluorescence of ATDC5 cells on normal, OA, and severe OA gels during control (dark blue) and 28 hr of 30 nM GSK101 (light blue) treatments. f-actin was used to draw ROI around the cells to measure TRPV4 fluorescence from max projections of z-stack images. Error Bars are ± SEM, * indicates p-value < 0.05.

Gene expression of anabolic and catabolic factors were analyzed to determine if decreasing substrate elasticity altered the phenotype of chondrocytes (Fig 5A). mRNA expression of *Sox-9*, a key chondrogenic transcription factor that is stimulated by TRPV4 activity, (Muramatsu et al., 2007) was not different across any stiffness. However, *aggrecan* and *col2* expression significantly decreased on the OA and severe OA gels when compared to the control normal gels. Activation of TRPV4 with GSK101 promoted an anabolic response by stimulating *aggrecan* and *col2* expression on the OA gels. However, substrate elasticity and TRPV4 stimulation had no effect on *collagen I* (*col1*) and *MMP-13* expression in our studies (Fig 5B). We also observed that both mRNA expression and total protein levels of TRPV4 decreased on the severe OA gels (Fig 5C).

**Figure 5.**
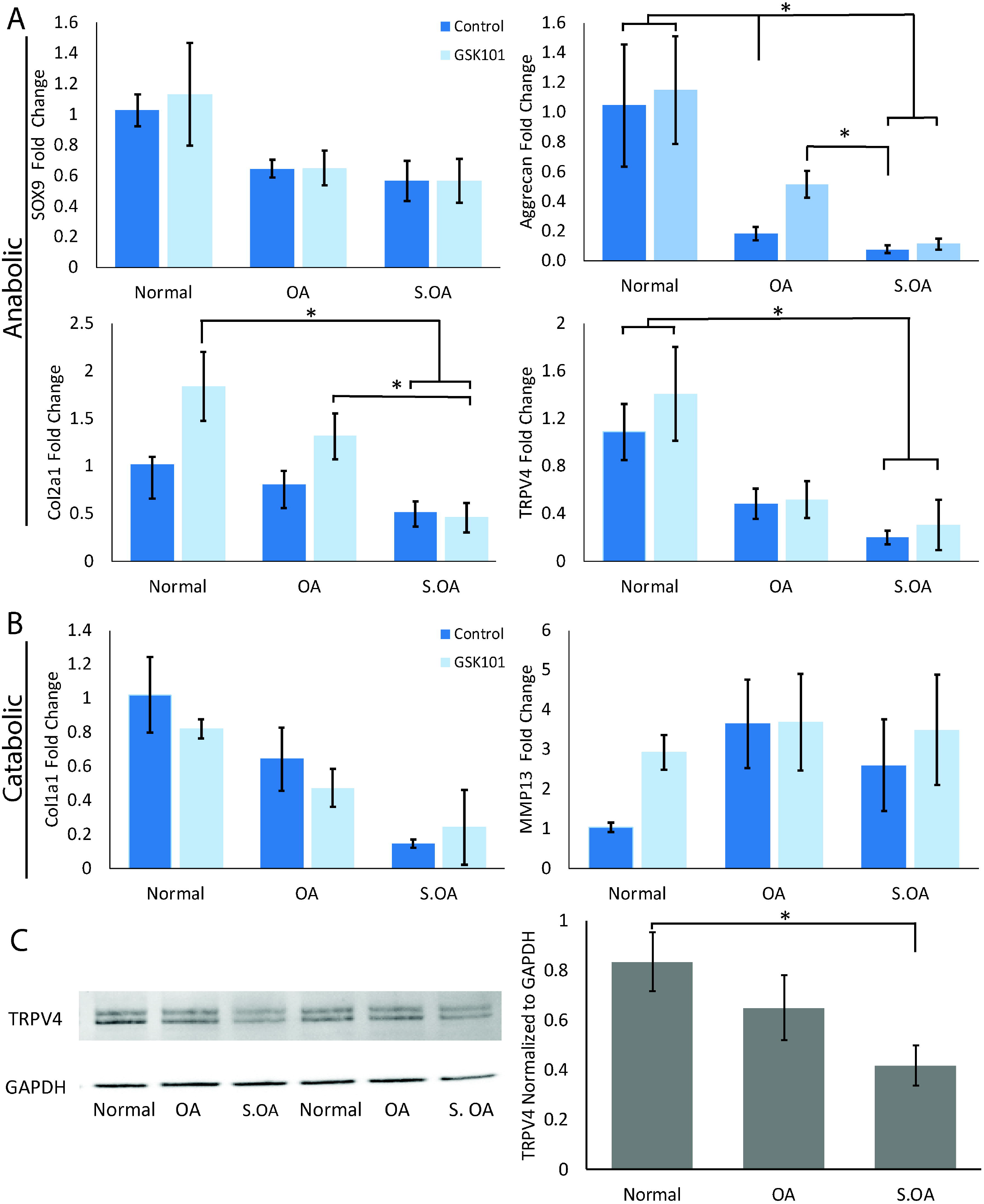
A) mRNA expression of anabolic factors (Sox-9, aggrecan, col2 and TRPV4) in ATDC5 cells on normal, OA and severe OA gels during control (dark blue) or 28 hr of 30 nM GSK101 (light blue) treatment. B) mRNA expression of catabolic chondrocyte factors, collagen I and MMP-13 in ATDC5 cells on normal, OA and severe OA gels during control (dark blue) or 28 hr of 30 nM GSK101 (light blue) treatment. Fold change was normalized to *rplp0* expression for each sample and then normalized to the normal control condition. C) Western blotting shows TRPV4 protein levels for ATDC5 cells on normal, OA, and severe OA gels during control treatment. The bar graph represents the normalized TRPV4 protein level to GAPDH for each sample. Error bars are ± SEM, * indicates significance with a p-value < 0.05.

## Discussion

During the progression of OA, the mechanical forces applied to chondrocytes change due to ECM degradation that decreases the compressive modulus of the tissue. Here, we investigated how TRPV4-mediated mechanotransduction changes as substrate elasticity decreases. We observed that TRPV4 regulates [Ca^2+^]_i_ signaling in response to HTS on normal and OA cartilage gels but not the severe OA gels. **(Fig 2 and Table 1)**. In chondrocytes embedded in agarose, TRPV4 transduces dynamic mechanical stimuli to promote synthesis of col2 and proteoglycans resulting in a concomitant increase in the Young’s modulus of the construct (O’Conor et al., 2014). While this suggests TRPV4 signaling is beneficial for cartilage homeostasis, knockout of TRPV4 attenuates age-related OA in mice (O’Conor et al., 2016). These conflicting results indicate that TRPV4 mediated mechanotransduction may be dependent on the biomechanical and biochemical status of cartilage. Here, we show that TRPV4 activity is regulated by the substrate stiffness, suggesting that the degradation of cartilage that accompanies the onset of OA exacerbates the progression of the disease.

Chondrocyte apoptosis is expected during OA, (Hashimoto et al., 1998), yet we found that 78% of the ATDC5 cells seeded on severe OA hydrogels were apoptotic. This, coupled with our [Ca^2+^]_i_ data **(Fig 3A)**, support a previous observation that [Ca^2+^]_i_ spikes prior to the induction of chondrocyte apoptosis (Huser and Davies, 2007). In our experiments, chondrocytes on severe OA gels had significantly higher basal [Ca^2+^]_i_ when compared to cells on the normal and OA conditions, which may be related to chondrocyte apoptosis during OA. Our data suggests that stimulating TRPV4 in chondrocytes on severe OA gels decreased the number of apoptotic cells to ~56% **(Fig 3A)**. However, TRPV4 stimulation had no effect on inhibiting the number of apoptotic cells using the Caspase 3/7 assay. These conflicting results may be related to the targets of the two assays. Propidium iodide intercalates with DNA during late stages of apoptosis or in necrotic cells. Meanwhile, Caspase 3/7 directly measures two executors of apoptosis (Brauchle et al., 2014). Thus, the conflicting results may mark the difference between apoptotic and necrotic cells.

The high incidence of apoptosis in our study conflicts with previous studies that used chondrocytes embedded in gels with similar moduli as our severe OA hydrogel did not observe similar levels of apoptosis (Mauck et al., 2000; Nicodemus and Bryant, 2008; O’Conor et al., 2014; Schuh et al., 2012; Villanueva et al., 2009). Chondrocytes embedded within hydrogels may improve cell viability and decrease apoptosis, potentially by forming a pericellular matrix that protects chondrocytes from mechanical stimuli and cell damage (Peters et al., 2011). During OA, the pericellular matrix degrades, increasing the magnitude of mechanical forces applied to chondrocytes, which may explain the observed increase in apoptotic cells in our studies (Zhang et, 2015).

We show that cells on the severe OA gels had significantly higher basal [Ca^2+^]_i_ compared to the other groups. This increase in basal calcium may activate calcineurin, a calcium activated phosphatase implicated in the progression of OA (Yoo et al., 2007). Chondrocytes within OA lesions express higher amounts of calcineurin and that inhibition of calcineurin prevents cartilage loss and decreases metalloproteases, IL-1, and nitric oxide (Van Der Windt et al., 2012; Yoo et al., 2007). NFATc2, is also upregulated in OA and inhibits proteoglycan synthesis while promoting matrix degradation (Van Der Windt et al., 2012). We postulate that the increase in basal [Ca^2+^]_i_ levels observed may activate calcineurin and NFATc2 signaling that influences the production of ECM proteins. Substrate stiffness did not alter the magnitude of the Ca^2+^ peaks, which contradicts previous reports that substrate stiffness influences the magnitude of Ca^2+^ oscillations (Kim et al., 2009; Zhang et al., 2020). However, Zhang et al. used freshly isolated chondrocytes from juvenile mice seeded onto collagen 1, which induces osteogenesis/hypertrophy in chondrocytes (Joyce et al., 1990). Juvenile and adult chondrocytes also exhibit differential spontaneous [Ca^2+^]_i_ oscillations, with adult chondrocytes having decreased amplitude of [Ca^2+^]_i_ peak and prolonged relaxation times compared to juvenile chondrocytes (Zhou et al., 2016). Thus, additional studies are necessary to determine if juvenile and adult chondrocytes similarly respond to substrate stiffness.

During the progression of OA, the balance between anabolic and catabolic factors shifts, with decreases in col2 and proteoglycan secretion, and concomitant increases in col1, inflammatory cytokines, and metalloproteases that degrade the ECM (Boileau et al., 2005; Ferrell et al., 2003; Goldring, 2001; Houard et al., 2013). Our results indicate that chondrocytes on softer substrates have large variations in col1 and MMP-13 production. Moreover, our data suggest that downregulation of aggrecan occurs in cells on OA and severe OA models when compared to cells on the normal gels **(Fig 5A)**. Treating cells with GSK101 abrogated this response to substrate elasticity. Furthermore, treating cells on the normal and OA conditions upregulated the expression of col2 when compared to the severe OA condition. While membrane bound TRPV4 is significantly higher on normal gels during GSK101 treatment when compared to the severe OA gels **(Fig 4A)**, TRPV4 mRNA and protein expression were downregulated on the severe OA gels compared to the normal gels, regardless of treatment **(Fig 5A and C)**. Previously, TRPV4 expression has been shown to increase during chondroprogenitor cell differentiation and in areas surrounding cartilage lesions during OA (Muramatsu et al., 2007; Soul et al., 2018). However, our results suggest that TRPV4 expression significantly decreases in cells on gels recapitulating severe OA.

In summary, this study confirms that TRPV4 mediates the response to HTS for chondrocytes for most conditions except when cells are on very soft substrates. We observed that TRPV4 stimulation could attenuate the amount of late apoptotic cells on severe OA gels but could not improve anabolic cytokine expression. We highlight that activation of TRPV4 improved the expression of *aggrecan* and *col2* on OA gels, indicating that TRPV4 may attenuate cartilage degradation during the early stages of OA.

## Acknowledgements

This work was supported in part by the National Institutes of Health U54GM104941 (RLD), R01HL133163 (JPG), R21CA214299 (JHS). J.H.S. was partially supported by a NSF Career Award (1751797). Microscopy access was supported by grants from the NIH (P20 GM103446), the National Science Foundation (IIA-1301765) and the State of Delaware. N.T. and C.J.F. were supported by the University of Delaware Graduate Scholar Award. Thank you members of the Gleghorn Lab for thoughtful discussions of this work.

