## Supplementary figures and images for "Extracellular Matrix Stiffness Alters TRPV4 Regulation in Chondrocytes"

### Supplemental Figure 1

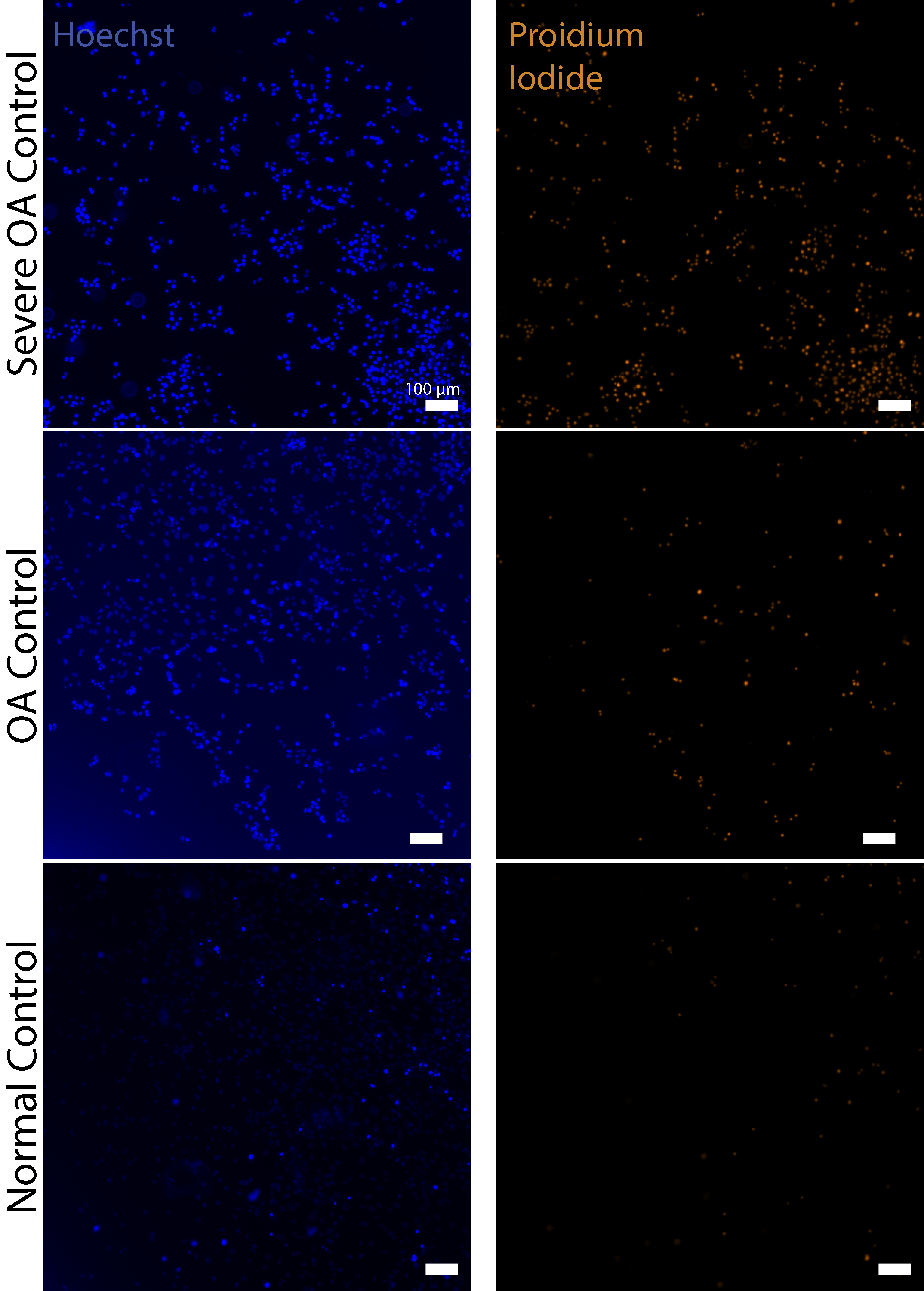

### Supplemental Figure 2

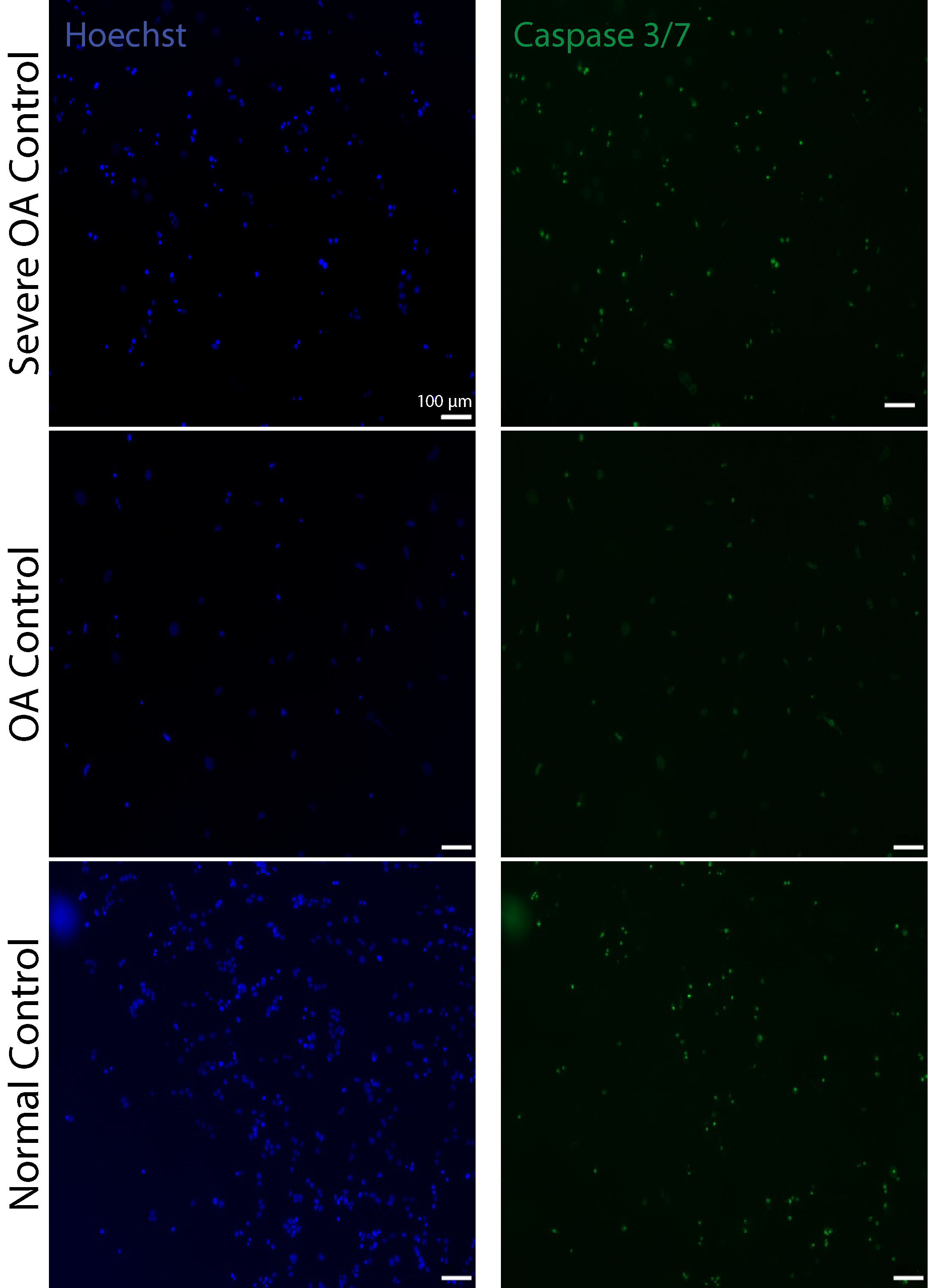

### Supplemental Figure 3

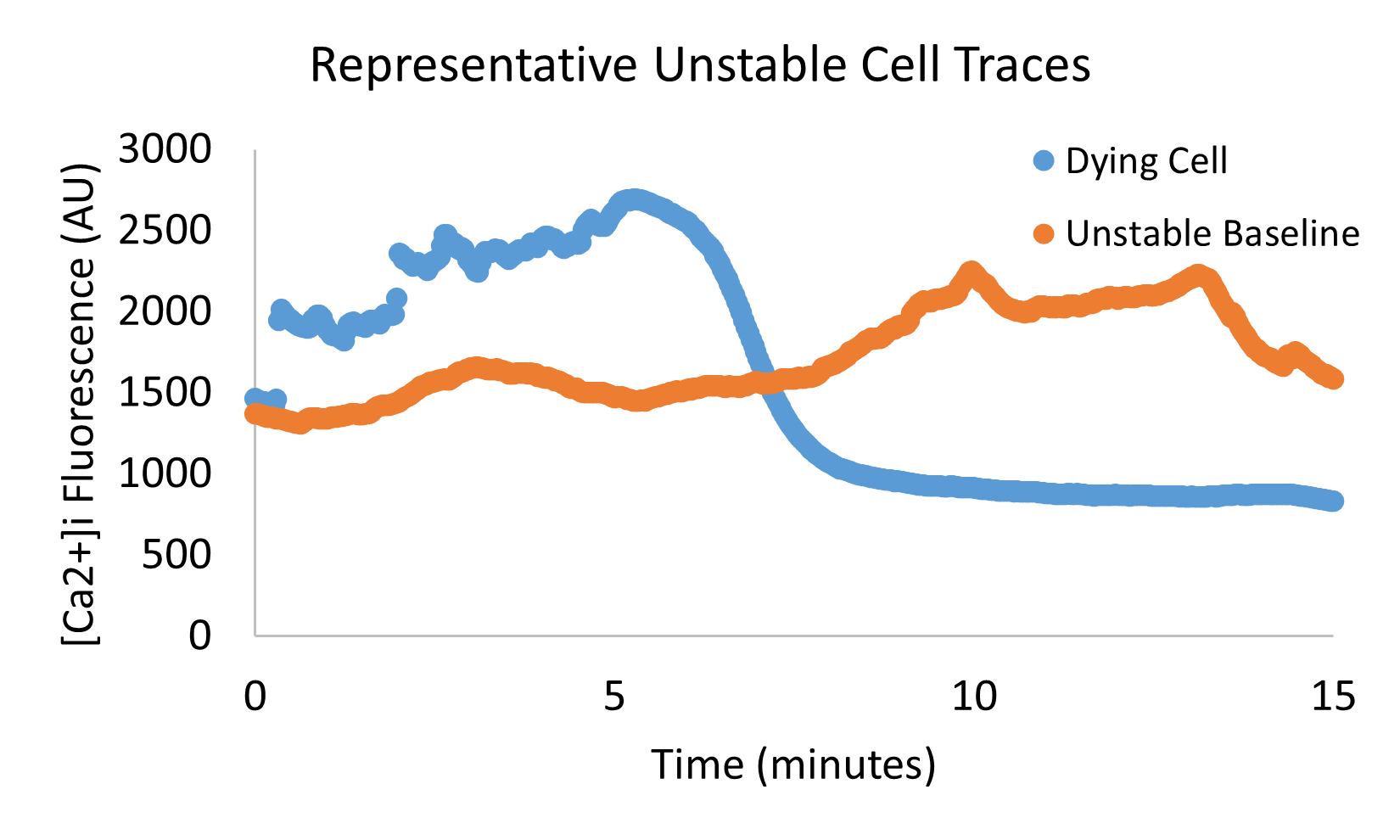

### Supplemental Figure 4

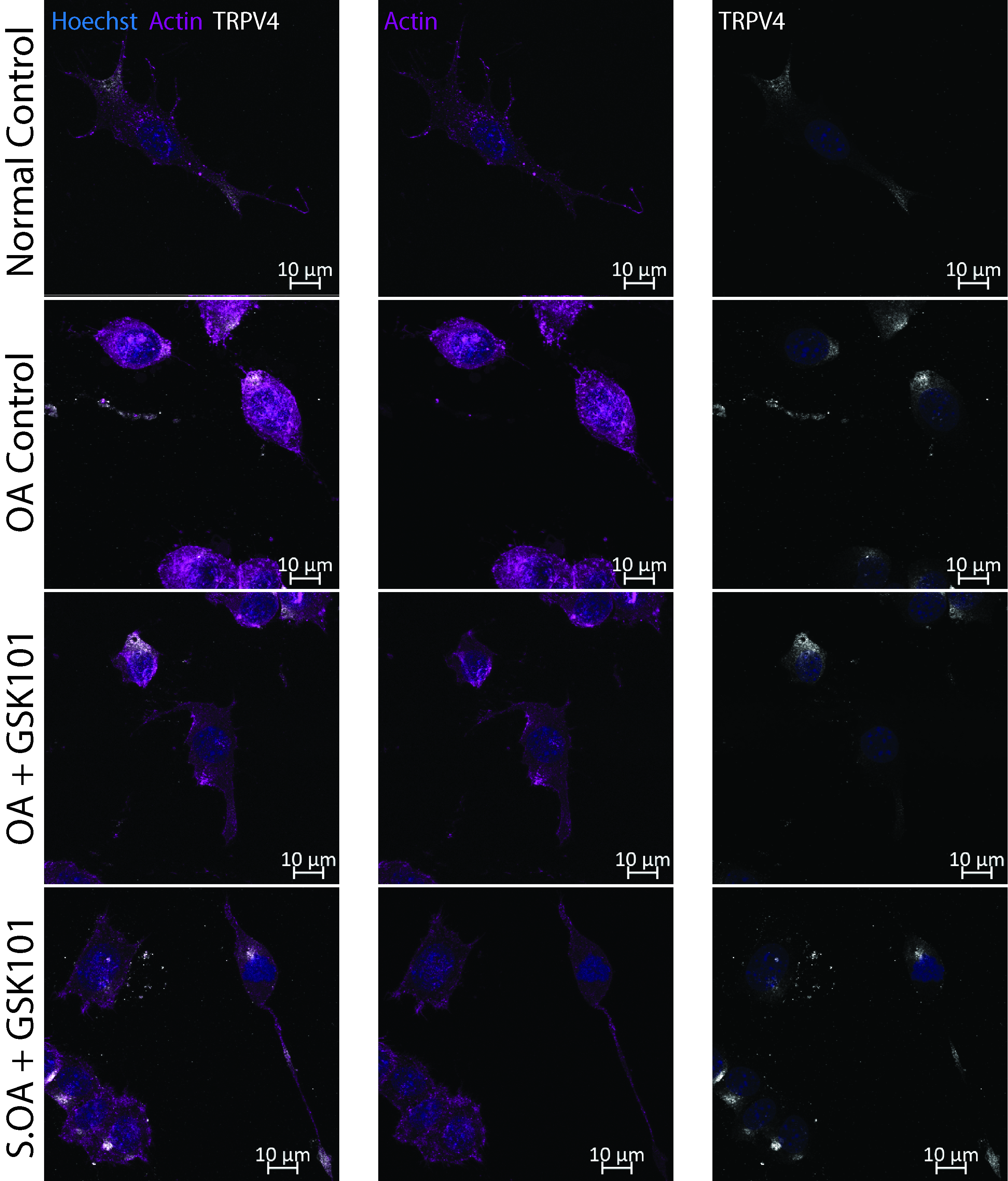
